## Supplemental information for "The evolution of antibiotic resistance is associated with collateral drug phenotypes in *Mycobacterium tuberculosis*"

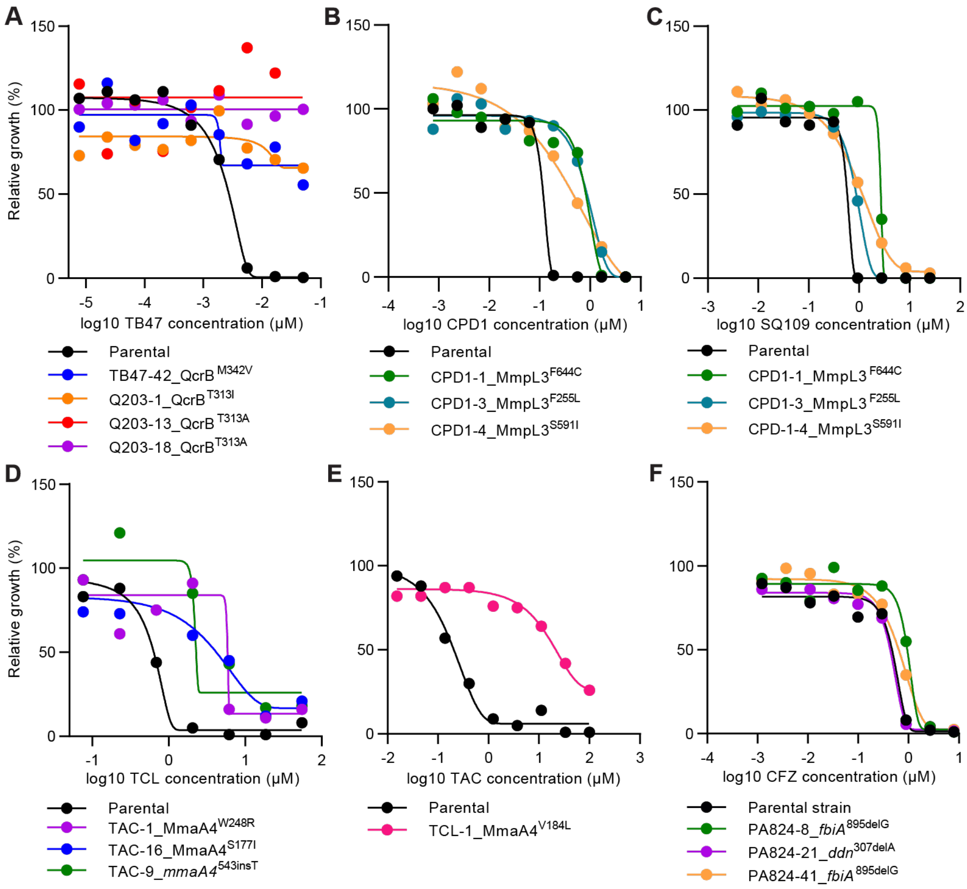


**Figure S1: Cross-resistance in drug-resistant variants of *M. tuberculosis*.** (A-F) examples of cross-resistance seen in dose-response curves for selected drug-resistant variants and the drug-susceptible parent against selected antibiotics. Strain names and candidate mutations are listed below each dose-response. Dose-response curves are the results of a single biological replicate from a representative experiment (n>2).

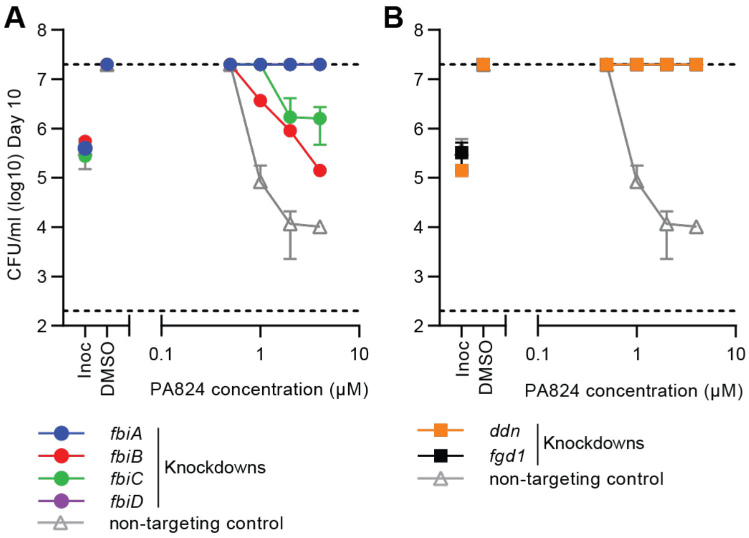


**Figure S2: Minimum bactericidal concentration assays for CRISPRi knockdown strains against PA824.** (A+B) Minimum bactericidal concentration assays for CRISPRi knockdown and non-targeting strains against PA824. CFUs were determined at day 0 and at day 10. Inoc = starting inoculum on day 0, DMSO = solvent control. Data is the mean ± SD of biological duplicates from a representative experiment (n>2). Dashed lines represent the upper and lower limits of detection.

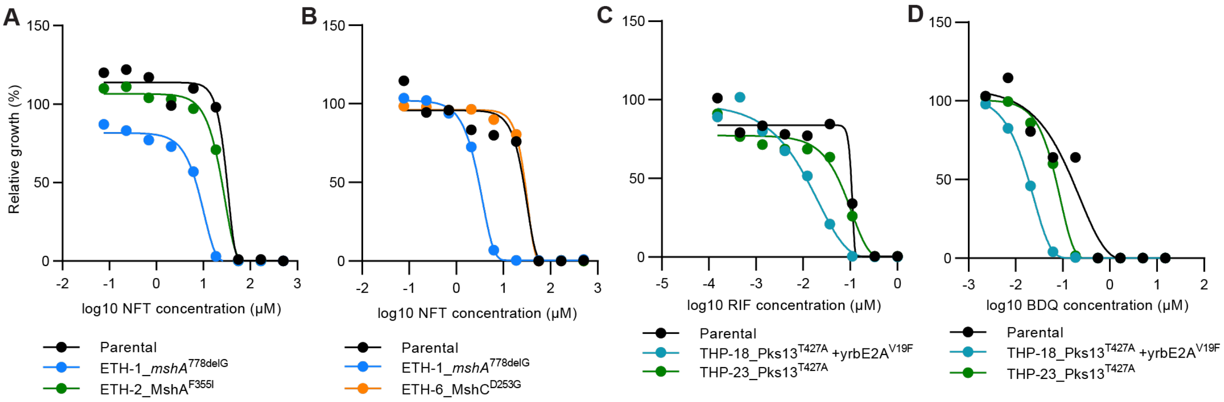


**Figure S3: Collateral sensitivity in drug-resistant variants of *M. tuberculosis*.** (A-D) examples of collateral sensitivity seen in dose-response curves for selected drug-resistant variants and the drug-susceptible parent against selected antibiotics. Strain names and candidate mutations are listed below each dose-response. Dose-response curves are the results of a single biological replicate from a representative experiment (n>2).

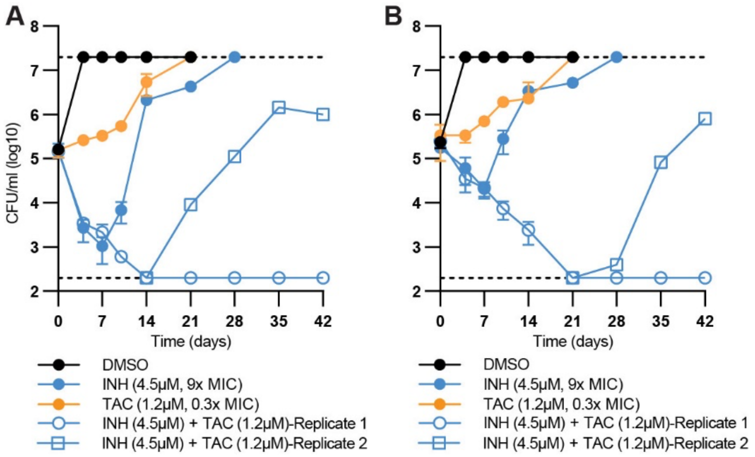


**Figure S4: Additional replicate experiments of INH + TAC combination studies.** (A+B) Drug-susceptible *M. tuberculosis* was incubated with either INH at 9x the MIC, below MIC concentrations (i.e., 0.3x MIC of the drug-susceptible parent) of TAC, or a combination of INH (9x MIC) and 0.3x MIC of TAC. CFUs were determined on the stated days. Data is the mean ± SD of biological duplicates from a representative experiment. Dashed lines represent the upper and lower limits of detection.

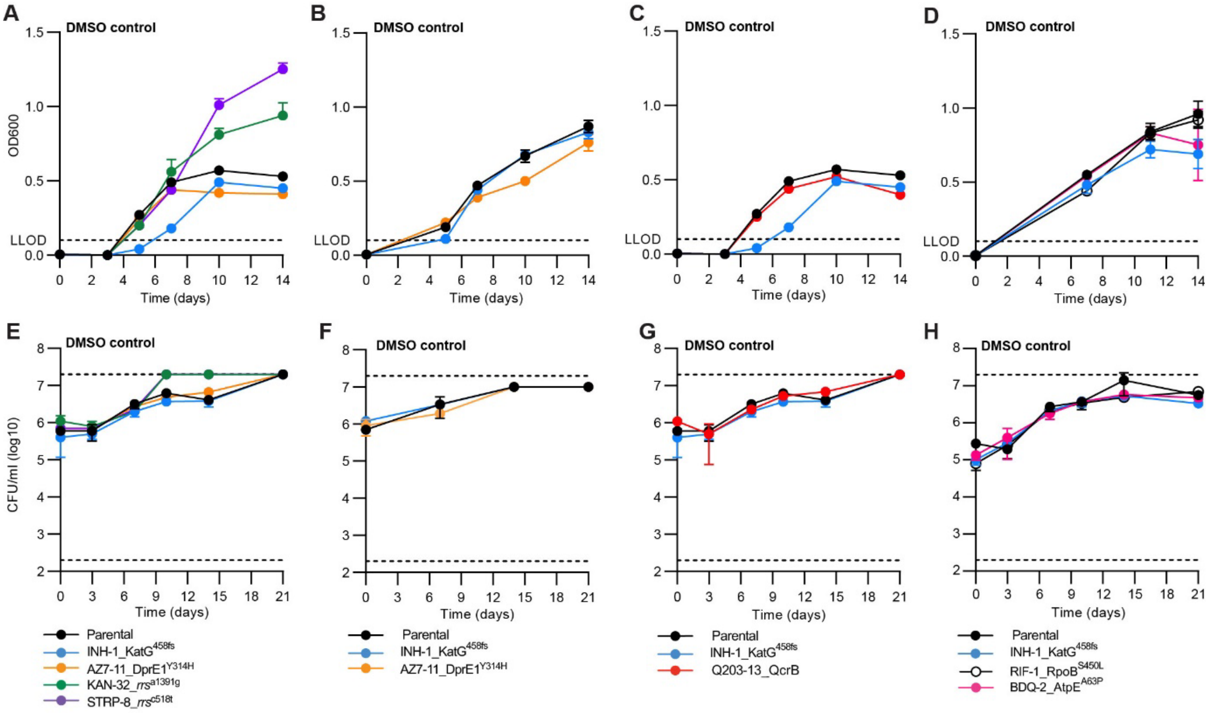


**Figure S5: DMSO control curves for relevant assays.** (A-D) Solvent control $OD_{600}$ curves for Figure 3E-H. (E-H) Solvent control CFU curves for Figure 4E-H. Dashed lines represent the upper and lower limits of detection.
